# Redefining Housekeeping Genes in the Context of Vertebrate Development

**DOI:** 10.1101/2025.11.12.687949

**Authors:** Alicia Lou, Juan F Poyatos, Monica Chagoyen

## Abstract

Housekeeping genes are typically defined as genes active in all adult tissues and essential for basic cellular functions. However, this view overlooks genes that are crucial during early development. Using zebrafish as a model system, we examined gene expression from the first hours of embryogenesis through adulthood. We found that many genes expressed throughout development remain broadly expressed in adults, but others do not; and some adult housekeeping genes are not active early on. This reveals that essential gene functions shift across life stages. Genes active throughout development are more likely to be evolutionarily ancient, shared with humans, and cause defects in many organs when disrupted. Yet some genes expressed only at specific developmental stages, especially around gastrulation, can be equally essential. This supports the developmental hourglass model, where mid-embryogenesis is the most conserved stage. Our results broaden the concept of housekeeping genes to include genes essential across time, not only across adult tissues.

## Introduction

Housekeeping (HK) genes are fundamental to maintaining basic cellular functions and are typically expressed at relatively constant levels across most cell types, developmental stages, and environmental conditions. In multicellular organisms, they are often contrasted with cell- or tissue-specific genes, whose differential expression underlies cellular identity and functional specialization.

The distinction between ubiquitous and specific genes is not only conceptually important but also highly practical. HK genes are widely used as internal controls for quantitative PCR and other normalization procedures (Zhang & Li, 2004). More broadly, they provide insight into core cellular processes essential for homeostasis, and serve as anchors for comparative and evolutionary analyses (Bergmiller et al., 2012; Joshi et al., 2022). Likewise, identifying cell- or tissue-specific genes is critical for annotating cell types in single-cell transcriptomics, revealing tissue-restricted disease mechanisms, and understanding the emergence of cellular diversity (Dvir et al., 2022; Kitsak et al., 2016; Li et al., 2025). Comparative studies between HK and tissue-specific genes have also deepened our understanding of transcriptional regulation and chromatin organization (Dall’Agnese & Young, 2023; Dejosez et al., 2023; Russo et al., 2018; Schug et al., 2005; She et al., 2009).

Yet, defining HK genes remains challenging. No single dataset can capture gene expression across all cell types, tissues, developmental stages, and environmental contexts. Most genome-wide efforts have therefore focused on adult tissues and cell lines, using technologies such as SAGE (Velculescu, V. et al., 1999), microarrays (Chang et al., 2011; De Jonge et al., 2007; Hsiao et al., 2001; Lee et al., 2007), expressed sequence tags (EST) (Zhu et al., 2008), and bulk tissue RNA sequencing [e.g., (Eisenberg & Levanon, 2013; Fagerberg et al., 2014; Hounkpe et al., 2021; Tung et al., 2024)]. These studies, largely centered on humans and mice, have yielded valuable HK gene sets but remain overwhelmingly adult-focused. A few multicellular species comparisons have broadened this view (Joshi et al., 2022), yet still predominantly rely on post-developmental tissues.

A developmental perspective, defining HK genes by temporal rather than spatial consistency, remains comparatively underexplored, despite its biological relevance. This approach emphasizes temporal consistency in gene expression during embryogenesis, as opposed to spatial uniformity across differentiated adult tissues. While a few studies have incorporated embryonic or fetal tissue alongside adult tissue data e.g., (Ramsköld et al., 2009; Warrington et al., 2000), or examined preimplantation embryos at the single-cell level (Lin et al., 2019), they have covered only limited differentiation phases. Notably, developmental gene expression has been also used to identify a short list of zebrafish housekeeping genes suitable for qPCR normalization, either in combination with adult tissue data and perturbation experiments (Xu et al., 2016) or concentrating on early stages of organismal growth (Y. Hu et al., 2016)

In this study, we take a systematic approach to HK gene definition from a developmental perspective using zebrafish (*Danio rerio*), a well-established vertebrate model. Using bulk RNA-seq data spanning key stages of embryogenesis (White et al., 2017), we classified genes into three temporal groups: ubiquitous (U) genes expressed across all stages, stage-specific (S) genes expressed in multiple but not all stages, and highly stage-specific (hS) genes restricted to narrow developmental windows. We then asked whether embryonic ubiquity predicts adult ubiquity, a cornerstone of classical HK gene definitions, by comparing our classification to a comprehensive adult tissue atlas (P. Hu et al., 2015). While there is substantial overlap, we also find notable mismatches, suggesting that essentiality and ubiquity are shaped by developmental timing as much as adult tissue identity.

To further interpret these gene classes, we characterize them in terms of their biological functions, anatomical pleiotropy, evolutionary age, and homology patterns across and within species. In doing so, we connect the concept of *developmental* HK genes to broader evolutionary frameworks such as the developmental hourglass model, which posits that mid-embryogenesis is the most conserved stage across species, marked by the expression of the oldest and most constrained genes. Our analysis provides evidence that developmental ubiquity captures this conserved core, while S and hS genes populate the more evolutionarily flexible early and late phases of development.

## Results

### Ubiquitous and stage-specific gene classes during vertebrate development

We introduced three gene categories based on an RNA-seq time-course dataset spanning 18 stages of zebrafish embryogenesis, from the zygote to the 5-day larval stage (White et al., 2017; Methods): *Ubiquitous* (U) genes (*n* = 8,579), expressed across all stages; *Specific* (S) genes (*n* = 7,551), expressed in ≥50% of stages; and *highly Specific* (hS) genes (*n* = 6,446), expressed in <50% of stages (Table S1, Figure 1A). An additional 1,066 genes did not meet classification thresholds (Figures S1A–B). As development progresses, the number of S and hS genes expressed increases, thus making the percentage of U genes expressed at each stage decline (Figure S1C). Despite this decline, U genes remain the largest and most stable fraction of the transcriptome, accounting for approximately 90% of total transcripts in early stages and about 71% in later stages (Figure S1D), with lower average individual gene expression variability in the earliest stages (till 16 hpf, when their average variability equals that of S genes), highlighting the stability of U genes throughout development, particularly in early embryogenesis (Figure S1E). It also highlights the low average variability in individual gene expression observed for hS genes at 4 and 5 dpf, the developmental stages at which their expression becomes increasingly important.

**Figure 1.**
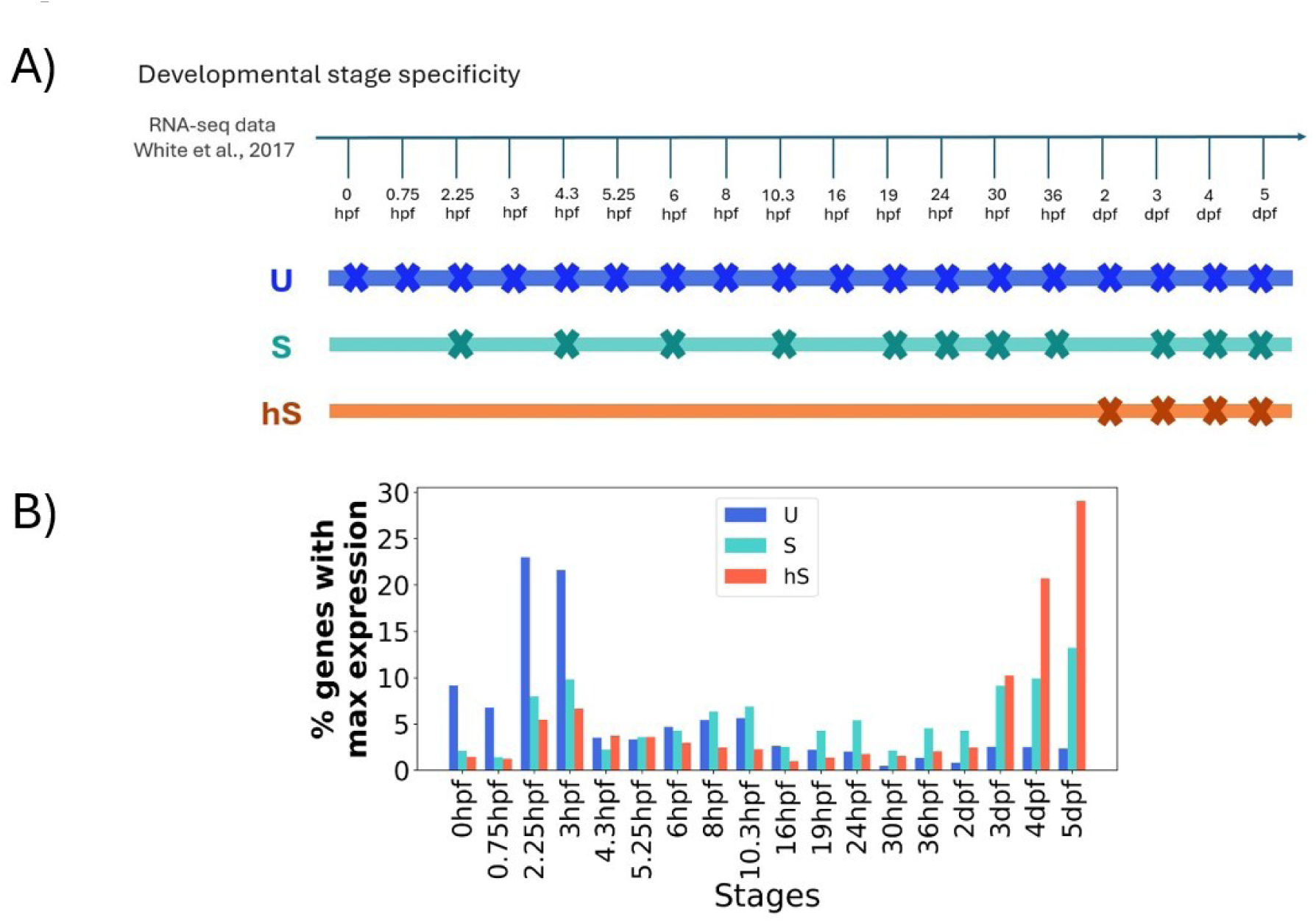
Gene classification based on developmental specificity. **A)** Developmental classification based on bulk RNA-seq embryos (White et al., 2017). The arrow represents all the developmental stages identified in the bulk data. Below, we show one example gene for each class. The X’s indicate that the gene is expressed at that stage (TPM > 1). The reason why the expression of the example hS gene is grouped in the latest stages is that hS genes begin to acquire transcriptional relevance between 30 hpf and 2 dpf (Fig. S1D). **B)** Distribution of genes by their stage of maximum expression (per developmental class). Each bar represents the percentage of genes in a given class that reach their highest expression at a specific developmental stage. For each gene class, the cumulative percentage across all stages adds up to 100% (hpf, hours post fertilization).

While our classification incorporates expression across *all* developmental stages, it is also informative to examine when each gene reaches its maximum expression. For each gene, we identified the developmental stage at which its expression peaks; this distribution helps distinguish the developmental classes (Figure 1B; Methods). U genes show their highest expression predominantly during zygotic genome activation (2.25-3 hpf), even though they account for most expression across early and mid-embryonic stages (Figure S1C-D). S genes exhibit a biphasic pattern, with peak expression occurring in both early and late stages. In contrast, and consistent with the progressive increase in overall transcription during development (Figure S1F), most hS genes reach their maximum expression at the latest developmental stages. Thus, beyond the differences in expression breadth, the distinction between S and hS genes also mirrors key developmental transitions.

### Developmental gene classes differentiate function and regulation

Nonetheless, it is reasonable to question how arbitrary this classification may be. To assess its biological validity, we first examined the functional associations of each group (Figure 2A, Table S2; Methods). As expected, U genes are enriched in core cellular functions such as biosynthesis and metabolic processes, RNA related processes, gene expression, and protein complex organization. By contrast, the two specific classes capture more specialized roles. S genes are enriched in transcriptional regulation and developmental processes, including cell differentiation, morphogenesis, and neurogenesis. hS genes, in turn, are associated with cell–cell and nervous system processes such as signaling, ion transport, and synaptic activity. These processes typically emerge by ∼20–30 hpf and continue developing into larval stages as neural circuitry matures.

**Figure 2.**
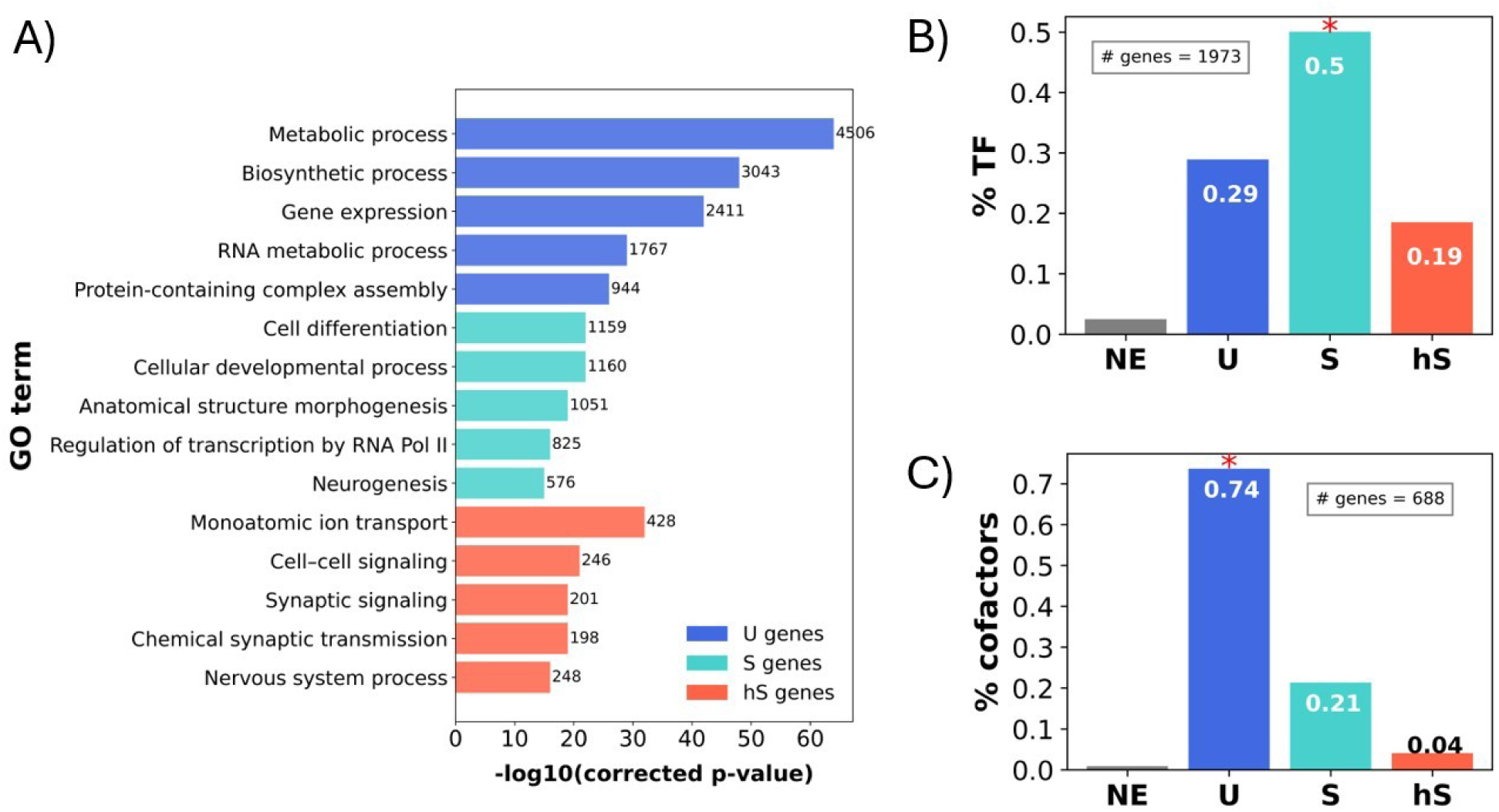
Developmental classes correspond to different functions and regulations. **A)** Enriched GO terms for each gene class: U (dark blue), S (light blue), hS (red). The y-axis lists GO terms selected from Table S2. Numbers next to each bar indicate how many genes are associated with that term, and enrichment significance is shown by the corrected p-value on the x-axis. **B)** Fraction of transcription factors (TFs) in each class relative to the total TFs in the dataset. **C)** Fraction of cofactors in each class relative to the total cofactors in the dataset. Asterisks above bars in (B) and (C) indicate classes significantly enriched based on a hypergeometric test (Methods, p < 0.0001).

We then compared transcription factors (TFs) and cofactors (including coactivators, corepressors, chromatin remodelers, mediator components, and TAFs), hypothesizing that their developmental expression patterns would reflect their distinct regulatory roles. Indeed, TFs remain largely inactive during the earliest stages of development and become expressed only after zygotic genome activation or during differentiation. This pattern is consistent with their predominant classification as S genes (50%, n=988), with limited association to U (29%, n=570) or hS (19%, n=366) categories (Figure 2B). Note that only the association with S genes remains statistically significant when testing for enrichment in each group (Methods, hypergeometric test, p-value=1.48e-68). This pattern corroborates findings in humans (Z. Hu & Gallo, 2010), mouse (Zhong et al., 2013), and other metazoan systems (Schep & Adryan, 2013). By contrast, cofactors are more evenly required throughout development, with the majority (74%, *n*=507; p=2.88e-91) falling within the U class, and only smaller and non-significative fractions in the S (21%, *n*=147) or hS (4%, *n*=28) groups (Figure 2C).

### Concordance between developmental and adult tissue gene expression classes

If our developmental classification captures meaningful biological features, how does it compare to the *standard* definition of HK genes in adulthood? To address this question, we analyze bulk RNA-seq data from eight major adult zebrafish tissues –brain, gill, heart, intestine, kidney, liver, muscle, and spleen (P. Hu et al., 2015) (Methods, Figure 3A). Genes were categorized into three groups as before but this time based on adult tissue expression patterns (Methods, Table S1): *ubiquitous tissue* (Ut) genes, *tissue-specific* (St) genes, and *highly tissue-specific* (hSt) genes (Figures S2A–B).

**Figure 3.**
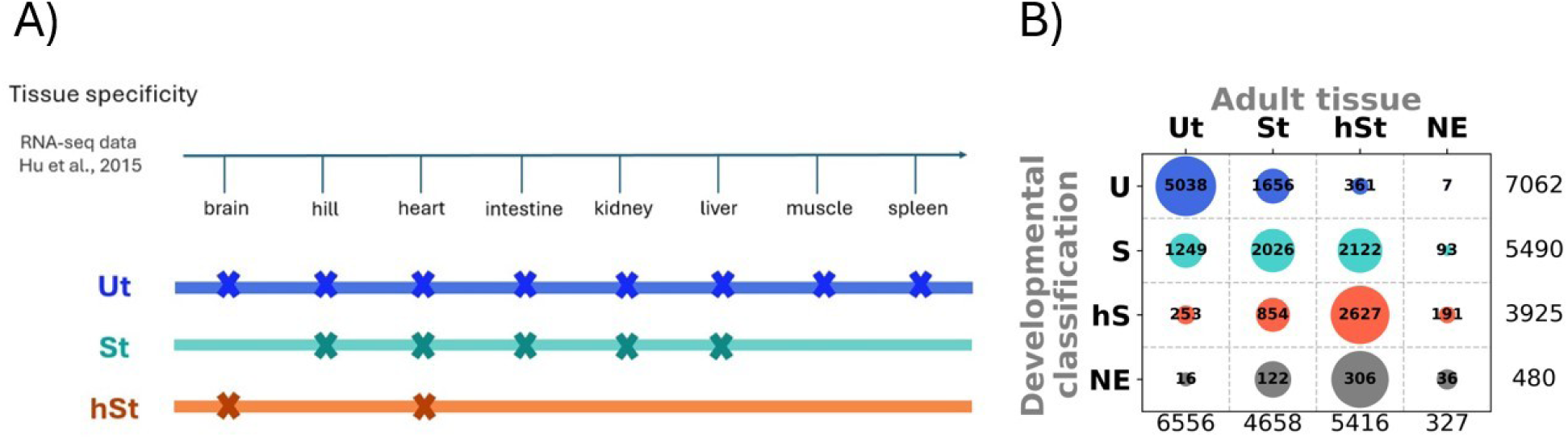
Gene classification based on tissue specificity. **A)** Adult tissue classification based on bulk RNA-seq data (P. Hu et al., 2015). The blue arrow depicts the set of tissues analyzed. Below, we present one example gene from each class, with X’s indicating the tissues in which that gene is expressed (TPM > 1). **B)** Comparison of developmental and tissue-based classifications. Points show the proportion of genes in each developmental class (U, S, hS) assigned to each tissue category (Ut, St, hSt), with point size indicating the row-wise percentage. While most genes retain their class, some fall outside the diagonal, indicating differences between the two schemes.

When compared with developmental expression profiles (Figure 3B, Table S1), approximately 71% of U genes retained ubiquitous expression in adult tissues (Ut type), indicating only partial conservation and suggesting post-embryonic, context-dependent regulation. These (U,Ut) genes are strongly enriched for core cellular processes, including RNA processing and translation, hallmarks of traditional HK function (Table S3; Methods). In contrast, (U,St) and (U,hSt) classes were enriched for cell cycle regulation, DNA repair, and microtubule-based processes, consistent with their peak activity during early developmental stages characterized by high proliferative demand.

S genes generally retained specific or highly specific expression in adult tissues. Specifically, (S,St) were enriched for functions in supramolecular fiber organization, enzyme regulation, and intracellular signaling. Moreover, (S,hSt) genes are associated with developmental processes, TF activity and cellular differentiation, including neurogenesis, peaking around 3 days post-fertilization (dpf) as brain structures mature. Intriguingly, a notable subset (∼23%) of genes with stage-specific developmental expression displayed ubiquitous expression in adult tissues (S,Ut). These genes were predominantly associated with mitochondrial localization and small molecule metabolic processes, reflecting the transition toward increased metabolic demand in later embryonic stages (Mendelsohn & Gitlin, 2008; Stackley et al., 2011).

Among hS genes, 67% remain highly specific in adult tissues (hSt). These genes are enriched for functions at the cell periphery, including plasma membrane localization, synaptic and cell-junction components, and ion channel activity. In addition, hS genes classified as St show enrichment in extracellular matrix organization, cell surface signaling, and immune response; functions broadly used across adult tissues but typically activated later in development. Notably, a small subset of hS genes becomes ubiquitously expressed in adulthood (hS,Ut), with roles in immune regulation and vitamin D metabolism, suggesting functions that are dispensable in early development but essential across tissues later in life.

Finally, expression level also varies across subclasses, with hSt genes showing significantly lower average expression than other subcategories (Figure S2C).

### Distinct properties of developmental gene classes

These analyses show that our developmental classification adds a temporal dimension to the *standard* HK framework, revealing distinctions not seen in adult tissues. This motivates examining how it relates to pleiotropy, gene family history, and evolutionary age.

#### Pleiotropy is not always coupled to developmental ubiquity of expression

For each gene, we quantified its organism-level phenotypic impact by measuring anatomical pleiotropy, defined as the number of tissues or structures reported in the Zebrafish Information Network as affected by its genetic or experimental perturbation (Methods; Supplement, Figure S3A). U genes display significantly higher anatomical pleiotropy than hS genes, but show no significant difference from S genes (Figure 4A). The proportion of genes with phenotypic annotations is similar for U and S genes (U: 2225/8579, ∼26%; S: 1635/7551, ∼22%), whereas hS genes have the lowest proportion (790/6446, ∼12%). No systematic differences were observed among tissue-based subclasses within each developmental class (U-Ut, etc., Figure S3B).

**Figure 4.**
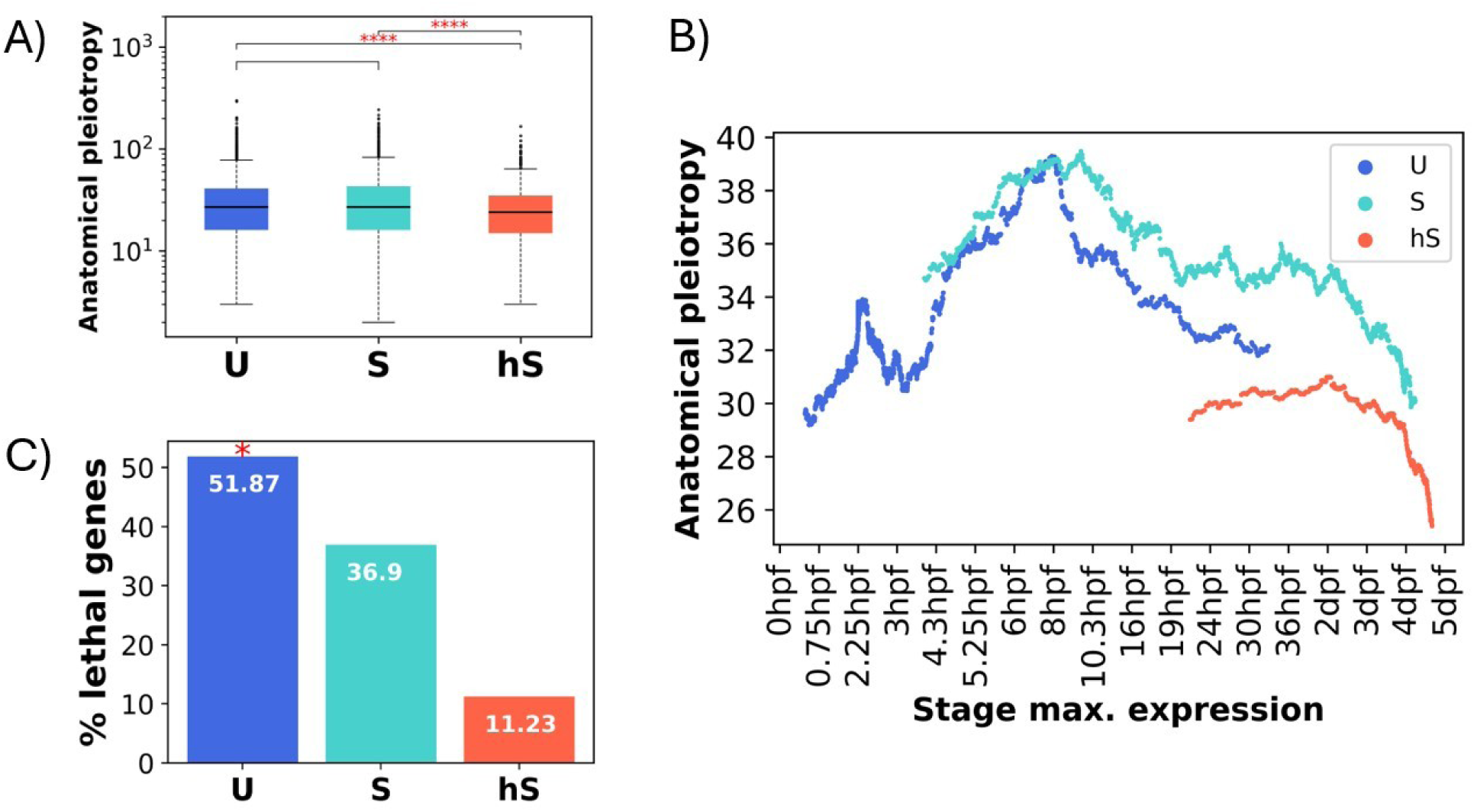
Anatomical pleiotropy analysis. **A)** Anatomical pleiotropy distribution among developmental classes. **B)** Average anatomical pleiotropy by stage of maximal expression, per gene class (sliding window=400, note that ∼26%, ∼22% and ∼12% of U, S, and hS genes, respectively have associated phenotypic effects). The apparent absence of S and hS genes at the earliest stages reflects missing anatomical annotations rather than absence of expression, as some reach maximal expression (Figure 1B) but likely lack associated phenotypic data. **C)** Fraction of genes with a ‘lethal’ phenotype in each class (U, S, hS), relative to all ‘lethal’ genes in the dataset. Asterisk indicates U class is significantly enriched based on a hypergeometric test (Methods, p < 0.0001).

Because the stage of peak expression is a defining feature of gene classes during development (Figure 1B), we assessed its relationship to gene pleiotropy using a sliding-window approach (Methods). Both U and S genes follow a similar trajectory: pleiotropy increases from the zygotic stage, peaks at 6–8 hours post-fertilization (shield/epiboly during gastrulation), and then declines, suggesting that the timing of peak expression is a major determinant of pleiotropy (Figure 4B).

More specifically, U genes are enriched in a broader set of general anatomical terms (Table S4; Fisher exact test: e.g., eye, head, liver) and are critical from gastrulation to early organ formation; their rapid decline in pleiotropy after the peak reflects simultaneous effects on multiple general structures. In contrast, S genes are enriched in fewer, more specific terms (e.g. portion of tissue, vein…) and sustain elevated pleiotropy over a longer developmental window, reflecting sequential effects on tissues and vascular structures. Finally, hS genes show relatively stable pleiotropy from 24 hpf to 3 dpf, followed by a decrease, and are consistently the least pleiotropic across all stages.

We also examined lethality, defined here as any genetic mutation that results in embryonic, larval, or adult death (an extreme manifestation of pleiotropy) and analyzed the distribution of 187 genes annotated as lethal. These genes are significantly enriched among U genes (97 of 8,579; hypergeometric test, p-val=9.0e-06), but show no significant enrichment in either S genes (69 of 7,551) or hS genes (21 of 6,446,) (Figure 4C). In addition, lethal U genes predominantly peak at 2.25 hpf, with additional peaks at 0 hpf, 3 hpf (zygotic genome activation), and 6 hpf (gastrulation onset) (Figure S3C). Lethal S genes (>10%) peaks at 8–10 hpf, stages associated with the highest pleiotropy in S genes and not prominent in the overall S gene expression profile. In contrast, lethal hS genes peak later, mainly at 4 dpf, with secondary peaks at 3 hpf (as observed for some hS genes in Figure 1B) and 30 hpf during organogenesis, when hS genes gain transcriptional relevance (Figure S1D).

#### Phyletic age patterns reflect developmental and functional constraints

Next, we evaluated phyletic age. Given the general concordance between our developmental classification and *standard* HK genes, we anticipated that U genes would also be evolutionarily older, like HKs are (Freilich et al., 2005), and indeed, this was the case (Figure 5A; Methods), consistent with the idea that U genes encode core cellular functions under strong evolutionary constraint. Significant age differences were likewise observed among subclasses defined by adult tissue expression profiles (Figure 5B, note the qualitative difference in the U-Ut class).

**Figure 5.**
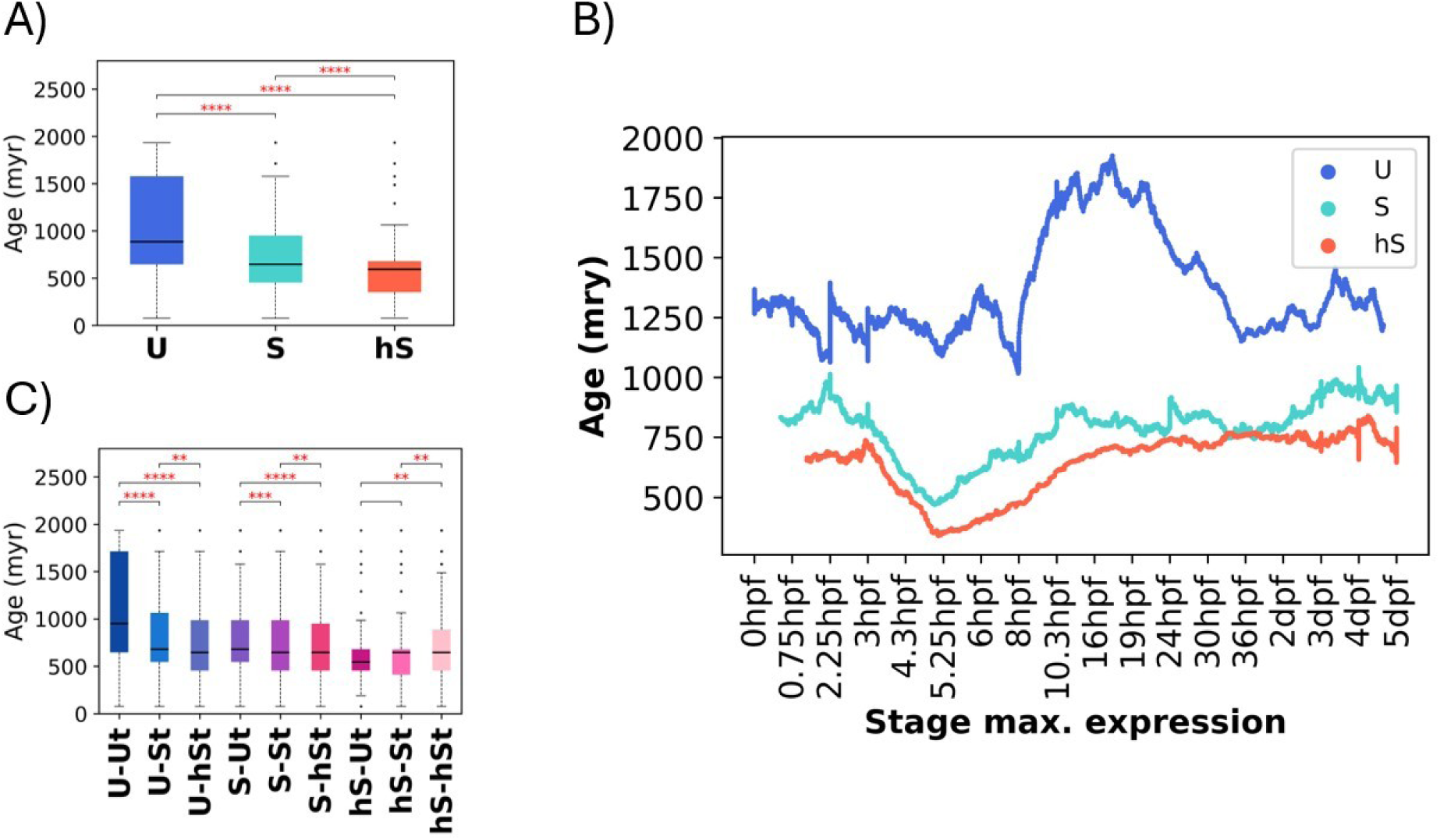
Analysis of phyletic age across development. **A)** Distribution of gene age per developmental class. All the distributions are significantly different; **** indicates a KS test p-value < 0.0001. We adjusted the y-axis limit and excluded outliers (genes with age >4290, labeled as 4290) to improve visualization of the distributions. **B)** Gene age vs. the maximal stage of expression in bulk data. Sliding window=300. **C)** Distribution of gene age per developmental and tissue-based subclass. Y-axis limits and outliers were handled as in A). ** indicates a KS test p-value < 0.01; *** p < 0.001; **** p < 0.0001. Lack of asterisks denotes a non-significant p value.

We again performed a sliding window analysis of age and peak expression per gene (Figure 5B; Methods). Note that our approach focuses on differences between gene classes across development rather than on global comparisons among stages, as in previous phylotranscriptomic studies employing the transcriptome age index (TAI) (Domazet-Lošo & Tautz, 2010; Ma & Zheng, 2024).

The youngest S and hS genes reach peak expression at approximately 5.25 hpf, immediately preceding gastrulation, a critical stage marking the onset of morphogenetic processes. Conversely, the most ancestral U genes tend to peak between 10.3 and 24 hpf, encompassing gastrulation, somitogenesis, neural tube formation, and the onset of organogenesis. This corroborates the hourglass model of development, which posits that phylogenetically ancient genes tend to be expressed during the conserved mid-embryonic phase (the “phylotypic” stage), whereas younger, lineage-specific genes are more often expressed at earlier or later stages (Irie & Kuratani, 2014).

We performed a functional enrichment analysis on the mentioned ancient U genes (phyletic age > 4,290 million years) with peak expression at stages 10.3, 16, 19, and 24 hpf, corresponding to the conserved mid-developmental phase (zebrafish phylotypic stage) (Schmidt & Starck, 2004; Table S5). Using the remaining U genes as a background, we identified significant enrichment for functions related to ribosomal structure and protein synthesis. These include processes such as ribosome biogenesis, ribonucleoprotein complex assembly, and translation (including initiation, elongation, and termination).

Moreover, evolutionarily young genes (S and hS <600 Ma) with peak expression at 5.25–6 hpf revealed distinct patterns (Table S5). S genes are enriched for nuclear processes, including transcription regulation, DNA binding, chromatin organization, and nucleic acid metabolism. hS genes share these functions but also show enrichment for immune response, receptor signaling, extracellular localization, and somite formation. Although young, these genes show nuclear and transcriptional functions, likely reflecting their integration into preexisting regulatory networks alongside older genes to control essential biological processes.

The fine-grained tissue subcategories identify further differences. U-Ut genes represent the most evolutionarily ancient subset (Figure 5C), with expression peaking between 10 and 19 hpf (Figure S4A). In contrast, U-St and U-hSt genes are significantly younger. Note that the U-hSt genes, developmentally ubiquitous but adult tissue–specific, are the youngest within the U class. Their expression peaks during blastulation and they are enriched in cell cycle–related functions, particularly those involving the cytoskeleton and microtubule dynamics (Table S3). This pattern is consistent with their role as genes required throughout development, when cell division is essential, but later restricted to adult tissues where cell proliferation remains especially active.

Interestingly, while the Ut subclass tends to include the oldest genes in both U and S categories (Figures S4A-B), this trend is reversed in the hS class (Figure S4C). The hS-Ut genes, despite being classified as adult-tissue ubiquitous, are among the youngest in the dataset, often younger than their hS-St and hS-hSt counterparts. They are enriched in the immune system process and response to external stimuli (Table S3), underlying the ubiquitous presence of immune cells across tissues. Altogether, this pattern indicates their broad incorporation into immune-related functions across tissues.

#### Gene family histories reveal the split between housekeeping and stage-specific roles

Finally, as the evolutionary age of a gene is closely associated with its family history, e.g., highly conserved genes typically possess numerous orthologs across species (Gabaldón & Koonin, 2013), we now investigate gene homology across species –focusing on orthology between zebrafish and humans– and within species –examining paralogous relationships that emerge from gene duplication events in zebrafish (Methods).

Consistent with their ancient origin, the majority of U genes have human orthologs (85% – n=7303; hypergeometric test, p-value=0.0), compared to 66% (n=4980) of S genes and 49% (n=3168) of hS genes (Figure 6A; Figure S5A to see the distribution of orthologs per gene subclasses), aligning with our earlier findings on gene evolutionary age.

**Figure 6.**
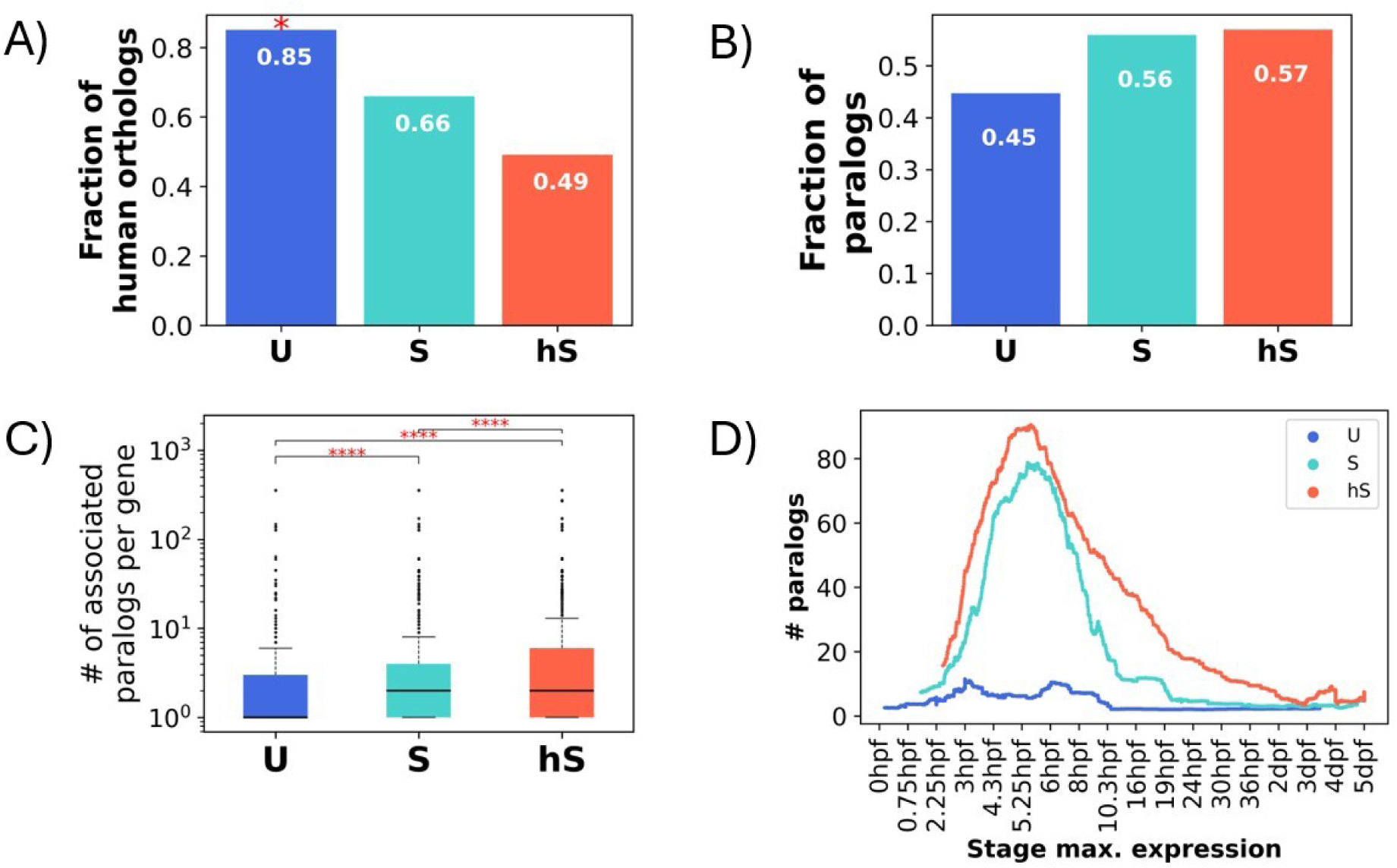
Homology analysis. **A)** Fraction of U, S, and hS genes with at least one human ortholog, relative to the total genes in each class. Asterisk indicates U class is significantly enriched based on a hypergeometric test (Methods, p < 0.0001). **B)** Fraction of U, S, and hS genes with at least one in-species paralog, relative to the total genes in each class. No class is significantly associated. **C)** Distribution of the number of paralogs per gene in each gene class. ** indicates a KS test p-value < 0.01; *** p < 0.001; **** p < 0.0001. **D)** Number of associated paralogs per gene vs. the maximal stage of expression in bulk data. Sliding window=400.

Moreover, paralogous genes, i.e., homologs within the same genome, drive functional innovation in multicellular organisms. Duplication followed by divergence allows them to adopt specialized roles, and prior studies show they are enriched in multicellular lineages and critical for expanding cell-type diversity (Padawer et al., 2012). Consistent with this, S and hS genes are more frequently associated with paralogs (S: 56% – n=4223; and hS: 57% – n=3672) than U genes (45% – n=3835); yet, no class is significantly enriched (Figure 6B; Figure S5B to see the distribution of paralogs per gene subclasses). Of note, the distribution of the number of paralogs per gene differs significantly across classes (Figure 6C).

In addition, we evaluated again how the number of paralogs varies across stages of peak expression using a sliding window analysis (Figure 6D). S and hS genes include genes with a high number of paralogs that reach maximum expression between 5.25 and 8 hpf, during gastrulation. After this point, paralog numbers drop earlier in S genes (reaching low levels by ∼10.3 hpf), whereas in hS genes the decline is more gradual, reaching similar values only by ∼3 dpf. In contrast, U genes show a consistently low number of paralogs throughout development, with only a modest increase during gastrulation. These patterns further support the hourglass model, highlighting gastrulation as a stage where genetic redundancy may be particularly relevant, and where innovation may take precedence over conservation.

Are there shifts between classes among paralogs? This may indicate regulatory divergence and the emergence of alternative developmental programs. Some pairs retain both genes within the same class (U–U: 32%, S–S: 51%, hS–hS: 41%), but the majority have at least one paralog assigned to a different class: 68% for U genes, 49% for S genes, and 59% for hS genes. These shifts are not random. Rather, paralogs often transition to the most adjacent class in terms of developmental breadth: U genes tend to have S-class paralogs, S genes most often pair with hS paralogs, and hS genes frequently have both S-class paralogs and paralogs that are not expressed during embryogenesis. (Table S6).

## Discussion

By considering developmental expression alongside adult profiles, we aim to broaden the conceptual framework of housekeeping (HK) genes. Rather than being restricted to those consistently expressed in mature tissues, HK genes may instead be better understood as those required across the life cycle, including the earliest stages of organismal development. This approach does not replace adult-focused analyses but complements them, providing a more comprehensive view of essential gene sets and their biological relevance.

We classified genes across zebrafish embryogenesis into ubiquitously expressed (U), specific (S), or highly specific (hS). This classification captures meaningful functional distinctions. U genes are enriched in core cellular activities such as RNA processing, translation, and macromolecular biosynthesis. S genes are enriched in transcriptional regulation, morphogenesis, and neurogenesis, while hS genes are associated with intercellular signaling, ion transport, synaptic function, and other specialized processes. As expected, essentiality is most enriched among U genes. Moreover, transcription factors are predominantly S-class, consistent with their roles in stage- and lineage-specific regulation, whereas transcriptional cofactors, often involved in basal transcription or chromatin remodeling, are largely U.

Comparison with adult tissue-based classifications revealed only partial overlap. Some developmentally ubiquitous genes become tissue-specific later in life, and some genes with restricted developmental expression become broadly expressed in adulthood. This highlights that temporal and spatial ubiquity capture different biological constraints. Developmental classifications thus reveal a layer of gene essentiality that is not recoverable from adult expression profiles alone.

We next examined functional and evolutionary properties associated with these developmental classes. First, anatomical pleiotropy –defined as the number of anatomical structures affected by perturbation– was highest among U genes and lowest among hS genes. Interestingly, S and U genes showed similar pleiotropy, despite differences in temporal breadth. Pleiotropy peaked during gastrulation (6–8 hpf), when early regulators influence multiple germ layers and future organ systems. These findings indicate that pleiotropy emerges either from sustained expression or from transient but developmentally pivotal activity.

Second, evolutionary analysis supports the developmental hourglass model, where mid-embryogenesis represents a highly conserved stage. Genes peaking during the zebrafish phylotypic period (16–19 hpf) (Schmidt & Starck, 2004) are the oldest, most conserved, and predominantly U. In contrast, genes peaking before gastrulation (∼5 hpf) are evolutionarily younger and enriched in the S and hS classes. This is consistent with transcriptome age index (TAI) studies showing that the phylotypic stage maximally expresses ancient genes (e.g., Domazet-Lošo & Tautz, 2010; Irie & Kuratani, 2014), whereas early and late stages recruit younger, lineage-specific genes. Our results further refine this view by showing that this evolutionary signal is embedded within developmental gene classes.

Third, contrary to the expectation that ancient genes accumulate more paralogs over time, U genes have fewer paralogs than S or hS genes. This likely reflects strong purifying selection, dosage sensitivity, or redundancy intolerance among core cellular genes (Rice et al., 2025). In contrast, S and hS genes –especially those involved in signaling, immune response, or developmental regulation– are more frequently duplicated, supporting the idea that gene duplication fuels innovation in regulatory and tissue-specific networks (Huminiecki & Heldin, 2010; Fernández & Gabaldón, 2020).

Together, these observations integrate with emerging refinements of the hourglass model. The phylotypic stage appears to function as a “developmental protocol layer”, highly constrained and dominated by ancient, ubiquitously required genes, while early and late stages allow greater evolutionary experimentation, recruitment of younger genes, and divergence between species. Our classification echoes this architecture: U genes largely populate the conserved core, while S and hS genes map onto the more flexible regulatory layers surrounding it.

More broadly, these results contribute to the evolving definition of HK genes. High-throughput transcriptomics has shown that many genes are weakly expressed in most tissues, undermining the classical definition of HK genes as those “expressed everywhere.” Instead, stability and indispensability across contexts, especially during early development, better capture housekeeping function. Our developmental classification aligns with this shift: U genes are not only broadly expressed but also evolutionarily ancient, pleiotropic, enriched in lethal phenotypes, and peak during phases of maximal developmental constraint.

Nevertheless, several limitations apply. Our dataset is based on bulk RNA-seq from whole embryos, which obscures cell-type specificity. Temporal coverage is limited to the first five days of development, excluding later stages and fully differentiated tissues. Adult datasets include only eight tissues, omitting others where important lineage-specific functions may emerge. These limitations should be considered when interpreting discrepancies between developmental and adult classifications.

Finally, our findings have implications for vertebrate evolution and biomedical research. U genes have human orthologs with essential roles and disease associations, reinforcing zebrafish as a model for conserved developmental processes. More importantly, the framework proposed here shows that gene essentiality arises through multiple routes –persistent ubiquity, temporally restricted regulation, or late specialization– and that these routes are embedded in evolutionary history.

In sum, the concept of HK genes expands from a static list defined in adult tissues to a dynamic, temporally informed framework. This integrated view bridges development, evolution, and gene regulation, and offers a foundation for future studies linking embryonic robustness, evolutionary constraint, and human disease.

## Methods

### Bulk RNA-seq data along development

The mRNA expression time-course data for zebrafish (*Danio rerio*) development were obtained by (White et al., 2017). This data includes transcriptomic profiles across 18 developmental stages, from the one-cell stage to five days post-fertilization. At each stage, transcriptomes were measured for five individual embryos. We downloaded the Transcripts Per Million (TPM) expression matrix from the Supplementary Materials of the publication. For each developmental stage, we calculated the mean TPM across the five embryos to obtain average expression levels (<TPM>). Because this averaging disrupts the fixed total of 1,000,000 TPM per sample, we performed an additional normalization step to correct for this discrepancy.

### Bulk RNA-seq data per tissue (adult zebrafish)

Tissue mRNA expression data for *Danio rerio* were obtained by (P. Hu et al., 2015). The dataset is available in the Gene Expression Omnibus (GEO) repository (www.ncbi.nlm.nih.gov/geo) under accession code GSE62221. Transcriptomic profiles were measured across eight tissues (brain, gill, heart, intestine, kidney, liver, muscle, and spleen) under three temperature conditions: 28°C, 18°C, and 10°C. The original expression matrix was reported in Reads Per Kilobase of transcript per Million mapped reads (RPKM). For our analysis, we transformed the data into Transcripts Per Million (TPM). Only data at 28°C were analyzed in this work.

### Gene Classification by Tissue Specificity and Comparison with Developmental Classes

Using bulk RNA-seq tissue data (see Tissue Data), we classified genes based on the number of tissues in which they were expressed (TPM > 1). We defined three categories: ubiquitous tissue (Ut) genes, expressed in all analyzed tissues; tissue-specific (St) genes, expressed in >50% of tissues; and highly tissue-specific (hSt) genes, expressed in <50% of tissues. We then intersected this tissue-based classification with the developmental gene classes (U, S, hS) and quantified the distribution of each developmental class across the Ut, St, and hSt categories.

### Data availability for Gene–Phenotype Associations and Lethality Annotations

Gene–phenotype association data were downloaded from ZFIN (The Zebrafish Information Network, https://zfin.org/; Data Reports, 26 Jan 2025), the primary database for genetic and genomic information on zebrafish (*Danio rerio*). The ontology organizes phenotypes using anatomical terms, specifying which structures or tissues are affected by genetic or experimental manipulations (see Supplement).

To quantify anatomical pleiotropy, we counted the total number of distinct anatomical terms associated with each gene, regardless of the developmental stage in which they appeared, noting that the number of terms increases with developmental complexity (Figure S3A).

Lethality information was also extracted from ZFIN. Genes annotated with the phenotype ‘lethal (sensu genetics)’ were retrieved, resulting in 329 ZFIN gene entries. After mapping to Ensembl identifiers, 190 genes were retained, of which 187 were present in our bulk developmental RNA-seq dataset.

### Data availability for the gene origin section

Gene age estimates were obtained from GenOrigin (http://genorigin.chenzxlab.cn/), a database that allows exploration of gene ages by species, evolutionary age, and gene ontology (Tong et al., 2021). Some genes were assigned an evolutionary age “>4290”. For data analysis, these genes were treated as having an evolutionary age of 4290.

### Data availability for the gene homology section

Orthologous gene pairs between zebrafish and human were obtained from the ZFIN database (https://zfin.org/downloads). Specifically, the orthology data were retrieved from the “Human and Zebrafish Orthology” CSV file available under the “Orthology Data” section of the Downloads page. Zebrafish paralogs were retrieved from Ensembl BioMart (https://www.ensembl.org/biomart/martview) by selecting the *Danio rerio* dataset and the “Paralogues” option under Homology attributes.

### Data availability for the transcriptional mechanism section

Zebrafish transcription factors and cofactors were retrieved from Animal TFDB4 (https://guolab.wchscu.cn/AnimalTFDB4/#/). From 2,546 transcription factors, we identified 1,973 in our data. From 782 cofactors, we identified 688.

### Adjusted Coefficient of Variation

The adjusted coefficient of variation (CV) was calculated following (Kolodziejczyk et al., 2015), to obtain a mean-independent CV. For each gene, we computed the CV at each stage using TPM values from the five embryos per stage and then transformed it as log₁₀(CV²). Genes were ordered by overall mean expression, and for each gene, we subtracted the median log₁₀(CV²) of 50 neighboring genes with similar expression. Positive values indicate that a gene is more variable than genes with comparable expression, while negative values indicate greater stability.

### Calculation of the stage of maximal expression

For each gene, we identified the developmental stage at which its expression reached the maximum value across all stages. Then, for each gene class, we calculated the percentage of genes showing maximal expression at each stage. This analysis yielded the distribution of maximal expression stages for each gene class (Figure 1B).

### Hypergeometric Test

We used the hypergeometric test to assess whether specific gene classes were significantly enriched in functional categories, such as transcription factors, cofactors, or essential genes. The background set consisted of all genes in the bulk dataset, while the gene sets of interest included all genes associated with our data. For each class (U, S, hS) and their respective subpopulations, the test calculates the probability of observing the overlap with a functional category by chance, using the hypergeometric distribution. This approach determines whether the representation of genes in each category is greater than expected randomly.

### Sliding window

To identify patterns between two biological variables, we applied a sliding window approach using Python’s Pandas library, specifically the rolling().mean() method.

Genes were first sorted by their stage of maximal expression, establishing the x-axis (developmental trajectory). The corresponding values of the y-axis (the gene-associated magnitude) were then reordered to match this sorted trajectory. A sliding window of size *sw* was applied to both axes to compute a rolling average. For example, the first computed average corresponds to values from x[0] to x[*sw*-1]. The next window includes values from x[1] to x[*sw*], then x[2] to x[*sw*+1], and so on. This process continues until the end of the array. Positions with fewer than sw values return NaN, ensuring all averages are computed from complete windows. This technique reduces noise and highlights underlying trends that may not be apparent in the raw data.

### Gene Ontology enrichment analysis

We downloaded Gene Ontology (GO) annotations from Gene Ontology Consortium (format-version: 1.2, data-version: releases/2025-02-06). We built three gene-GO term association matrices one for each aspect of the ontology (molecular function, cell component and biological process), by expanding direct annotations to ancestor terms in the GO hierarchy. GO term enrichment was assessed for each gene subset (U, S, hS) and their derived subsets (e.g., U-Ut, U-St) using Fisher’s exact test (fisher_exact function from scipy.stats), comparing the respective gene set against the complementary set comprising all remaining genes in the dataset. We corrected p-values within each GO aspect for multiple testing using False Discovery Rate (FDR).

## Supplementary material Tables

**Supplement.** Supplementary analyses and figures S1–S5.

**Table S1.** List of genes classified across development using RNA-seq data obtained by White *et al*. (2017) and across tissues using RNA-seq data obtained by Hu *et al*. (2015).

**Table S2.** Gene Ontology (GO) enrichment of the three gene classes across development: U, S, and hS.

**Table S3.** GO enrichment of gene subsets: U-Ut, U-St, U-hSt, S-Ut, S-St, S-hSt, hS-Ut, hS-St, and hS-hSt.

**Table S4.** Enrichment of anatomical terms from ZFIN for U, S, and hS genes.

**Table S5.** GO enrichment analyses of gene subsets defined by maximal expression stage and phyletic age. Subsets of ancient U genes (phyletic age > 4,290 million years) with peak expression at stages 10.3, 16, 19, and 24 hpf, and subsets of young S and hS genes (phyletic age < 600 million years) with peak expression at stages 5.25 and 6 hpf were included.

**Table S6.** Classification analysis of paralog gene pairs, evaluating whether both genes in each pair are classified within the same developmental class or in different classes.

## Code availability

Github: https://github.com/ali4lou/Redefining-Housekeeping-Genes

